# Serotonin in the orbitofrontal cortex enhances cognitive flexibility

**DOI:** 10.1101/2023.03.09.531880

**Authors:** Jung Ho Hyun, Patrick Hannan, Hideki Iwamoto, Randy D. Blakely, Hyung-Bae Kwon

## Abstract

Cognitive flexibility is a brain’s ability to switch between different rules or action plans depending on the context. However, cellular level understanding of cognitive flexibility have been largely unexplored. We probed a specific serotonergic pathway from dorsal raphe nuclei (DRN) to the orbitofrontal cortex (OFC) while animals are performing reversal learning task. We found that serotonin release from DRN to the OFC promotes reversal learning. A long-range connection between these two brain regions was confirmed anatomically and functionally. We further show that spatiotemporally precise serotonergic action directly enhances the excitability of OFC neurons and offers enhanced spike probability of OFC network. Serotonergic action facilitated the induction of synaptic plasticity by enhancing Ca^2+^ influx at dendritic spines in the OFC. Thus, our findings suggest that a key signature of flexibility is the formation of choice specific ensembles via serotonin-dependent synaptic plasticity.

## Introduction

Behavioral flexibility allows animals to make efficient decisions in a given environment. In order to respond adaptively and generate proper behaviors in a fast-changing environment, we must be able to update new information quickly to the demands of the stimuli and ready to learn new one when contingency changes. Also, deficits in this executive function have been associated with a variety of brain disorders^1,2^. Thus, flexibility is a key cognitive feature of the brain, but circuits that control flexibility and the neurobiological mechanisms by which flexibility is encoded are unknown.

Previous studies reported that the OFC is essential for enabling one to flexibly switch behaviors when encountering unexpected outcomes. Electrophysiological recording experiments show that neurons in the OFC encode the value of an external environment^3-8^. Furthermore, lesions in the OFC area in human and non-human primates have led to deficits in choice behavior^9-13^. Reversal learning paradigms have served as reliable tests of cognitive flexibility across species; moreover, OFC neurons appear to play an important role in reversal learning^2,14-18^. Given that reversal learning measures behavioral flexibility when reward-related contingencies are reversed, researchers have evaluated the capability of reversal learning in order to understand mechanisms of cognitive flexibility.

The most well-characterized cellular substrate underlying cognitive flexibility might be the serotonin system. When serotonin was selectively depleted in the OFC area, animals showed cognitive inflexibility and impaired reversal learning^19,20^. Several other studies have demonstrated the importance of serotonin in reversal learning and cognitive flexibility^21-30^. Thus, there is indirect evidence that the circuit pathway from serotonergic neurons in the DRN to OFC is important for cognitive flexibility, but general unifying principles underlying cognitive flexibility, especially at cellular resolution, have not been proposed. A handful of research has hinted at a link between DRN and OFC^31-33^, and a recent study began to visualize the anatomical connection between these two brain areas^34^. Here, we investigated the circuit functions underlying cognitive flexibility by targeting the pathways from DRN serotonergic inputs to the OFC and by monitoring individual neuronal activities while animals perform RL tasks.

We performed experiments across a broad range of approaches, from testing synaptic plasticity at single dendritic spines to monitoring the population activity of OFC neurons. To demonstrate the necessity of circuits, we optically and pharmacologically perturbed a local brain area, and also examined behavioral causality by targeting specific cell-neural connections. By doing so, we uncovered the cellular principles of RL, which may be a general feature of cognitive flexibility in a mammalian brain.

## Results

### Activation of DRN Serotonergic Neurons Promotes Reversal Learning

We used a two-choice lever pressing behavior as a RL paradigm. In brief, our behavioral task in an operant chamber comprised of two phases, normal associative learning and RL sessions (Fig. 1a). During the RL period, water reward was provided only when mice pressed the active lever (left or right). After 9 rewards, suddenly, the active lever became inactive and the inactive one became active. This change in reward contingency should lead to animals figuring out the rule change and coming up with a new solution. Because we did not deliver any cue for the rule switch, animals normally kept pressing the incorrect lever that was used to yield a water reward. As a result, the total number of incorrect lever presses remained high on the first day of reversal, but the ratio of incorrect-to-correct lever presses (reversal ratio) gradually decreased over time (Fig. 1b). This result indicates that mice are capable of disengaging from a previously learned rule and then trying to find an alternative solution.

**Fig. 1.**
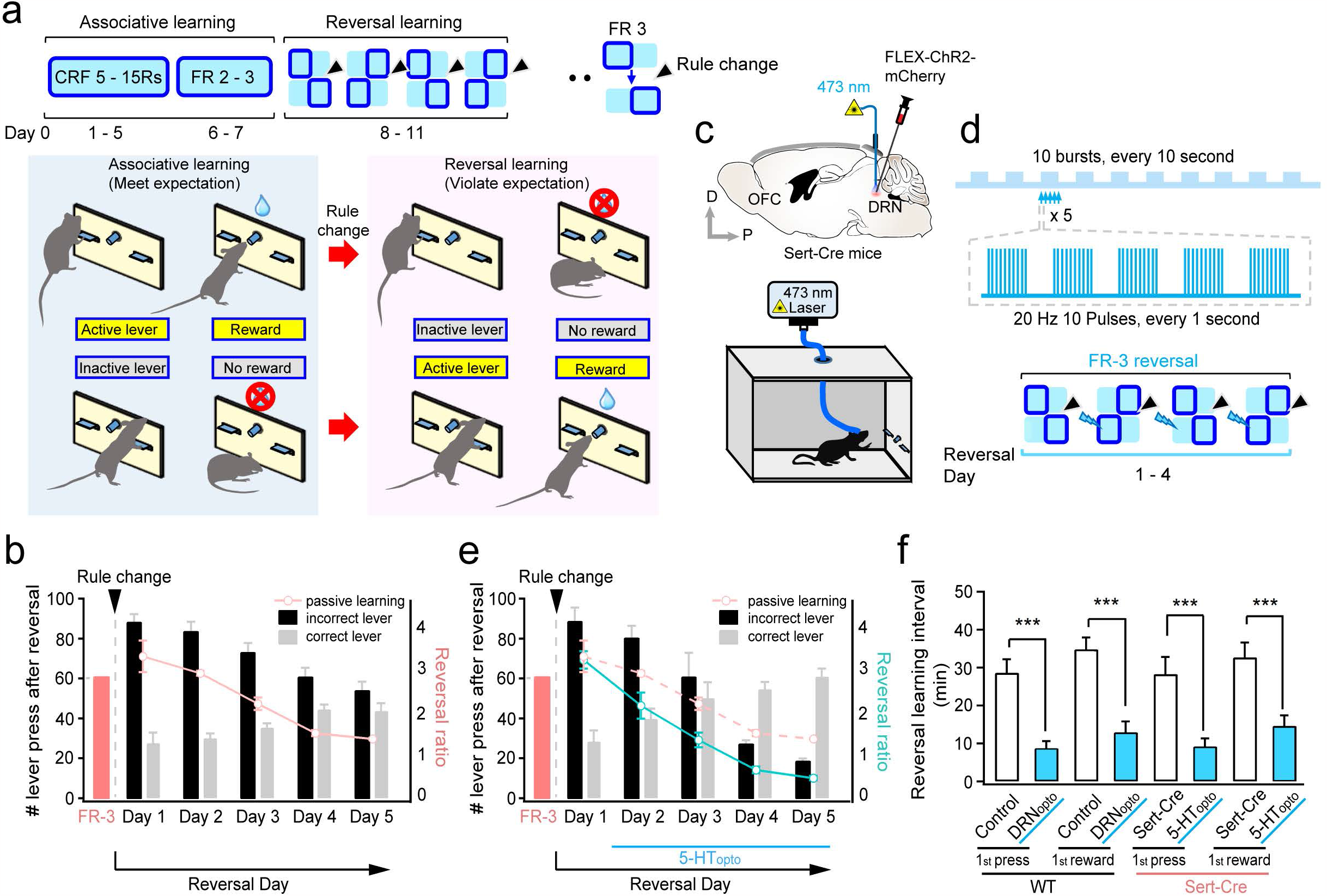
Serotonin improves reversal learning. **a**, A procedure of behavioral training plan (top). A schematic figure for reversal learning task (bottom). After the rule switch, active lever becomes inactive lever and *vice versa*. **b**, A summary graph of correct and incorrect lever pressing ratio change during the reversal learning phase. **c**, A schematic figure illustrating places for virus injection and fiber optic implantation. AAV-FLEX-ChR2-mCherry was injected into the DRN of Sert-Cre mouse. **d**, Protocols for blue light illumination. During the period of rule switch, blue light was repetitively delivered to activate serotonin neurons. Blue box denotes an active side lever. Angled arrowhead denotes the time point of rule change. **e**, A summary graph of reversal ratio changes. When blue light was illuminated, the ratio decreased faster, indicating reversal learning was improved (Passive learning: 5 animals, 5-HT_opto_: 7 animals (one-way repeated-measures (1-way RM)-ANOVA, day: *F*_(2.2,19.5)_ = 43.1, p < 0.005; 5-HT_opto_ effect: *F*_(1,9)_ = 26.2, p < 0.005). **f**, A summary graph of reversal learning interval (WT, 1^st^ press: control: 28.7 ± 3.6 min; DRN_opto_: 8.8 ± 1.8 min, n = 14, p < 0.005; WT, 1^st^ reward: control: 34.8 ± 3.1 min; DRN_opto_: 13.0 ± 2.8 min; n = 14, p < 0.005; Sert-Cre, 1^st^ press: control: 28.3 ± 4.5 min; 5-HT_opto_: 9.3 ± 2.1 min, n = 10, p < 0.005; Sert-Cre, 1^st^ reward: control: 32.7 ± 3.9 min; 5-HT_opto_: 14.7 ± 2.8 min; n = 10, p < 0.005). For all graphs, *** indicates p < 0.005, and Error bars indicate s.e.m.

To determine whether 5-HT facilitates RL, we injected adeno associated virus (AAV)-Flex-ChR2-mCherry into the DRN of Sert-Cre mice (Fig. 1c). Approximately 3-4 weeks after virus injection, the water-restricted mice were trained for normal associative learning to get water rewards. After rule change, we delivered repetitive high-frequency blue light pulses (10 pulses at 20 Hz, 5 times every 1 sec, repeated 10 times every 10 sec) to the DRN area during the reversal phase to activate serotonergic neurons (Fig. 1d). Triggering serotonin neuronal firing by the optogenetic activation of 5-HT neurons (5-HT_opto_) accelerated the speed of reversal, decreasing the reversal ratio faster compared to the control group without blue light illumination (Fig. 1e). The average time interval taken to press the other side of lever and get the first reward after the rule change was also shortened (Fig. 1f). To make sure whether blue light was sufficient to trigger action potentials from serotonergic neurons, we verified cell firing via a whole-cell current clamp recording from acute brain slices (Supplementary Fig. 1a).

### Anatomical and Functional Verification of DRN Serotonergic Projections to the OFC

We also verified the anatomical long-range projection from the DRN to the OFC. Retrograde beads (Supplementary Fig. 1b) or AAVrg-pCAG-FLEX-tdTomato was injected into the OFC of either wild-type (WT) or Sert-Cre mice, and the axonal projections from the DRN to the OFC or the presence of the OFC-projecting neurons in the DRN were clearly visualized (Fig. 2a-d). To confirm the functional release of 5-HT, we performed a fast-scan cyclic voltammetry (FSCV) recording by putting a carbon fiber microelectrode in the OFC (Fig. 2e). Upon blue light illumination, 5-HT release was detected in a dose-dependent manner (Fig. 2f,g). Thus, these anatomical mapping and optogenetic experiments showed that DRN-OFC circuits play an important role in RL.

**Fig. 2.**
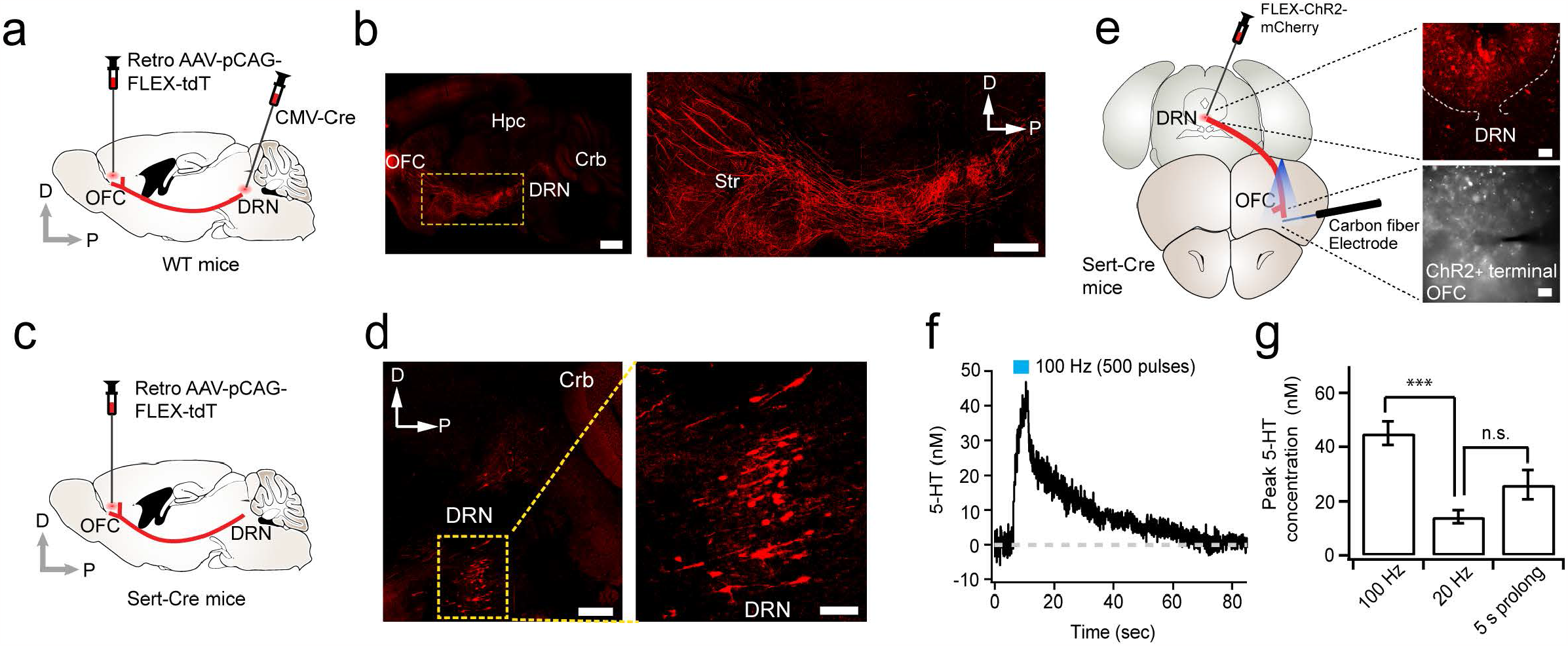
Anatomical and functional verification of long-range projection from the DRN to the OFC. **a**, Experimental scheme for Cre-dependent retrograde AAV and CMV-Cre injections into OFC and DRN of WT mice. **b**, Representative images of whole brain sagittal section visualized intact axonal projection from the DRN to the OFC. Scale bars indicate 1 mm (left) and 200 μm (right). **c**, Experimental scheme for Cre-dependent retrograde AAV injection to Sert-Cre mice. **d**, Representative sagittal images of selectively labeled serotonin neurons projecting to OFC. Scale bars indicate 100 μm (left) and 50 μm (right). (OFC, orbitofrontal cortex; DRN, dorsal raphe nucleus; Hpc, hippocampus; Crb, cerebellum; Str, striatum). **e**, Experimental scheme for fast scan cyclic voltammetry recording (left). Representative images depict ChR2+ neurons or terminals from DRN or OFC area, respectively. Scale bars, 100 μm (top) and 10 μm (bottom). f, Measurement of 5-HT in the OFC upon blue light illumination. **g**, Frequency- and duration-dependent 5-HT release (45.1 ± 4.4 nM for 100 Hz, n = 10, p < 0.005 compare to 20 Hz; 14.3 ± 2.4 nM for 20 Hz, n = 4, n.s. compare to 5 s; 26.3 ± 5.5 nM for 5 s, n = 4). For all graphs, *** indicates p < 0.005, and Error bars indicate s.e.m.

### Necessity and Sufficiency of DRN-OFC Activity for Reversal Learning

When triggering action potentials from serotonergic neurons in the DRN by ChR2 activation, 5-HT release should not be confined to the OFC area. To determine the specific role of 5-HT constrained to the OFC area, we injected retrograde AAV-floxed ChR2-mCherry in the OFC of Sert-Cre mice (Fig. 3a). Then, we implanted optic fibers in the DRN to activate only a subset of neurons (Fig. 3a; Supplementary Fig. 2a-d). Blue light illumination during the reversal period accelerated the speed of RL, as indicated by a faster reduction of the reversal ratio (Fig. 3a). To ensure that the observed RL change was indeed mediated through 5-HT in the OFC, we expressed ChR2 selectively in 5-HT neurons and delivered blue light directly on the axonal terminal of serotonergic fibers in the OFC (Fig. 3b). This manipulation significantly improved RL, confirming that indeed 5-HT action in the OFC mediates these flexible switching behaviors. Direct activation of 5-HT neurons by ChR2 might cause 5-HT release beyond physiological levels. To determine whether we can suppress RL by inhibiting 5-HT release during the RL task, we injected a retrograde virus expressing inhibitory designer receptors exclusively activated by designer drugs (DREADDs) (Retro-AAV-hSyn-DIO-hM4Di-mCherry) bilaterally into the OFC of Sert-Cre mice (Fig. 3c). After the virus was fully expressed (4∼5 weeks), we activated the DREADD receptors by using a synthetic ligand of DREADD, clozapine-N-oxide (CNO)^35^. CNO injection significantly reduced the reversal ratio (Fig. 3d) and subsequently prolonged reversal time interval (Fig. 3e). These changes were not due to a deficit in normal associative learning, because normal learning (pressing the correct lever) before the rule change was not affected by CNO (Supplementary Fig. 3g,h). The following day, CNO and fluoxetine were co-injected to see whether augmenting the amount of 5-HT can restore normal RL. Because fluoxetine blocks 5-HT re-uptake, released 5-HT stays longer in the brain^36^. Co-administration of CNO and fluoxetine fully restored the normal RL from the same animals that showed impaired RL on the previous day (Fig. 3d,e). To confirm whether retrograde hM4Di receptors are expressed enough to suppress the activity of DRN neurons, we performed whole-cell recordings from mCherry-expressing neurons in the DRN and applied CNO to brain slices. We injected a depolarizing current to evoke action potentials and found that CNO application successfully inhibited the firing of neurons (Supplementary Fig. 3a-f). These results showed that local release of 5-HT in the OFC is necessary for RL, and up- and down-regulation of the activity of 5-HT neurons have bidirectional effects on RL.

**Fig. 3.**
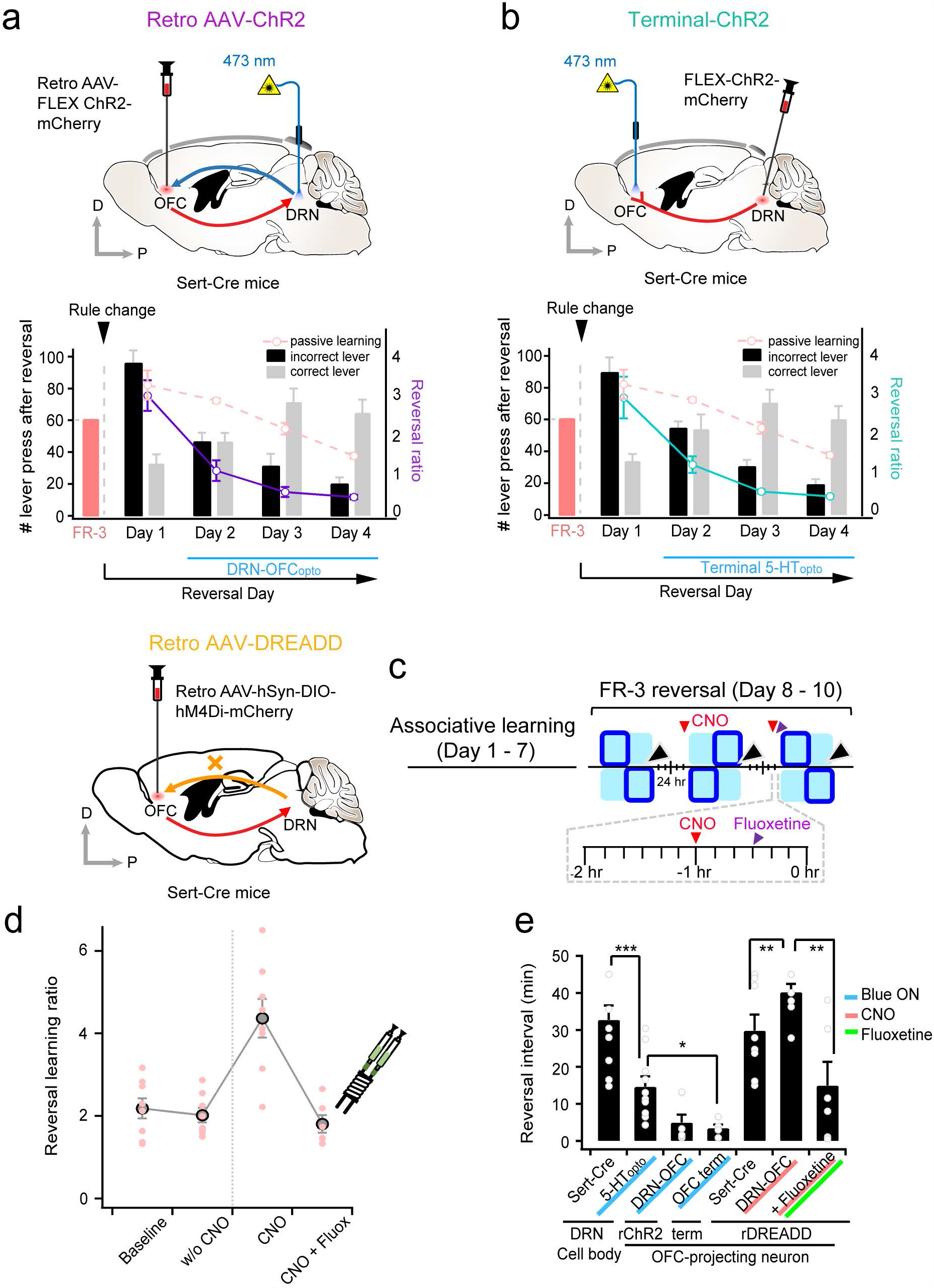
Specific serotonergic pathway from the DRN to the OFC is necessary for reversal learning. **a**, Retrograde AAV expressing floxed ChR2 was injected to the OFC of Sert-Cre mice and the optic fiber was implanted to the DRN in order to activate a group of 5-HT positive neurons that are projecting to the OFC (top). A summary graph representing the number of lever press after the reversal phase (5 animals) (bottom). The ratio of incorrect lever to correct lever was significantly reduced when serotonergic neurons were activated. **b**, Direct axonal terminal activation of 5-HT neurons in the OFC improved reversal learning. A schematic diagram of virus injection and optic fiber implantation (top) and the summary graph of reversal learning behavioral task (bottom) (1-way RM-ANOVA, day: *F*_(1.2,13.3)_ = 52.2, p < 0.005; 5-HT_opto_ effect: *F*_(2,11)_ = 15.8, p < 0.005; Dunnett’s T3 post-hoc test, control vs retro AAV-ChR2, p < 0.005; control vs terminal ChR2, p < 0.005; AAV-ChR2 vs terminal ChR2, p = 1.00). **c**, Experimental timeline for chemogenetic inhibition and the rescue effect by intracranial fluoxetine infusion on reversal learning. **d**, Schematic cartoon for surgery (upper) and the summary plot of reversal learning ratio changes (8 animals, Baseline: 2.2 ± 0.2, w/o CNO: 2.0 ± 0.2, n.s. compare to baseline, CNO: 4.4 ± 0.5, p < 0.005 compare to w/o CNO, CNO + fluoxetine: 1.8 ± 0.2, p < 0.005 compare to CNO). Note that, the action of 5-HT in the OFC mediated flexible update of behaviors. **e**, A summary bar graph of reversal intervals in various conditions as labeled (DRN cell body, Sert-Cre: 32.7 ± 3.9 min; 5-HT_opto_: 14.7 ± 2.8 min, p < 0.005 compare to Sert-Cre; rChR2: 5.0 ± 2.1 min; OFC term: 3.5 ± 0.9 min, p < 0.05 compare to 5-HT_opto_; rDREADD, Sert-Cre: 29.8 ± 4.3 min; CNO: 40.2 ± 2.2 min, p < 0.01, compare to Sert-Cre; Fluoxetine: 15.0 ± 6.4 min, p < 0.01 compare to CNO). *, ** and *** indicate p < 0.05, p < 0.01 and p < 0.005, respectively. Error bars indicate s.e.m.

### Serotonin Alters Internal Brain States in the OFC

To determine the cellular mechanisms underlying RL, we examined the extent to which 5-HT alters cellular properties and how these alterations impact RL. We monitored neuronal excitability, one of the effects of 5-HT observed in other brain areas^37,38^. We injected AAV-floxed ChR2-mCherry into the DRN of Sert-Cre mice (Fig. 4a). Then, we illuminated blue light directly into the OFC to trigger 5-HT release exclusively in the OFC. In a current-clamp mode, we injected depolarizing currents to layer 5 pyramidal neurons in the OFC and examined how 5-HT influences the magnitude of depolarization. Blue light illumination depolarized neurons, resulting in more action potentials (Fig. 4b). This change was due to the depolarization of resting membrane potentials (Fig. 4c). To examine more physiologically-relevant 5-HT action in cell firing, we tested the relationship between excitatory postsynaptic potentials (EPSP) to spike (E-S coupling). We used bipolar electrode to stimulate afferent fibers to the target neuron, and gradually increased the stimulation intensity to evoke EPSPs (Fig. 4d). We adjusted the stimulation strength to the level where action potentials are not initiated. After recording a stable baseline, we turned on the blue light and maintained the same stimulation to determine the effect of blue light on E-S coupling. As predicted, cells were depolarized and the number of action potentials were significantly increased in the presence of blue light (Fig. 4e; Supplementary Fig. 4a-e). 5-HT-induced inward current was also confirmed by a voltage-clamp recording mode (Fig. 4f), and the magnitude and decay kinetics of inward currents were enhanced in the presence of fluoxetine (10 μM) (Supplementary Fig. 5a-c). This effect was blocked by a 5-HT_2C_ antagonist, SB242084, but not by a 5-HT_1A_ antagonist, WAY 100635, indicating that 5-HT_2C_ receptor activation-mediated depolarization. These results also allowed us to exclude a possible involvement of glutamatergic effect on this 5-HT-induced inward current due to co-release of 5-HT and glutamate from DRN (Fig. 4g)^39^.

**Fig. 4.**
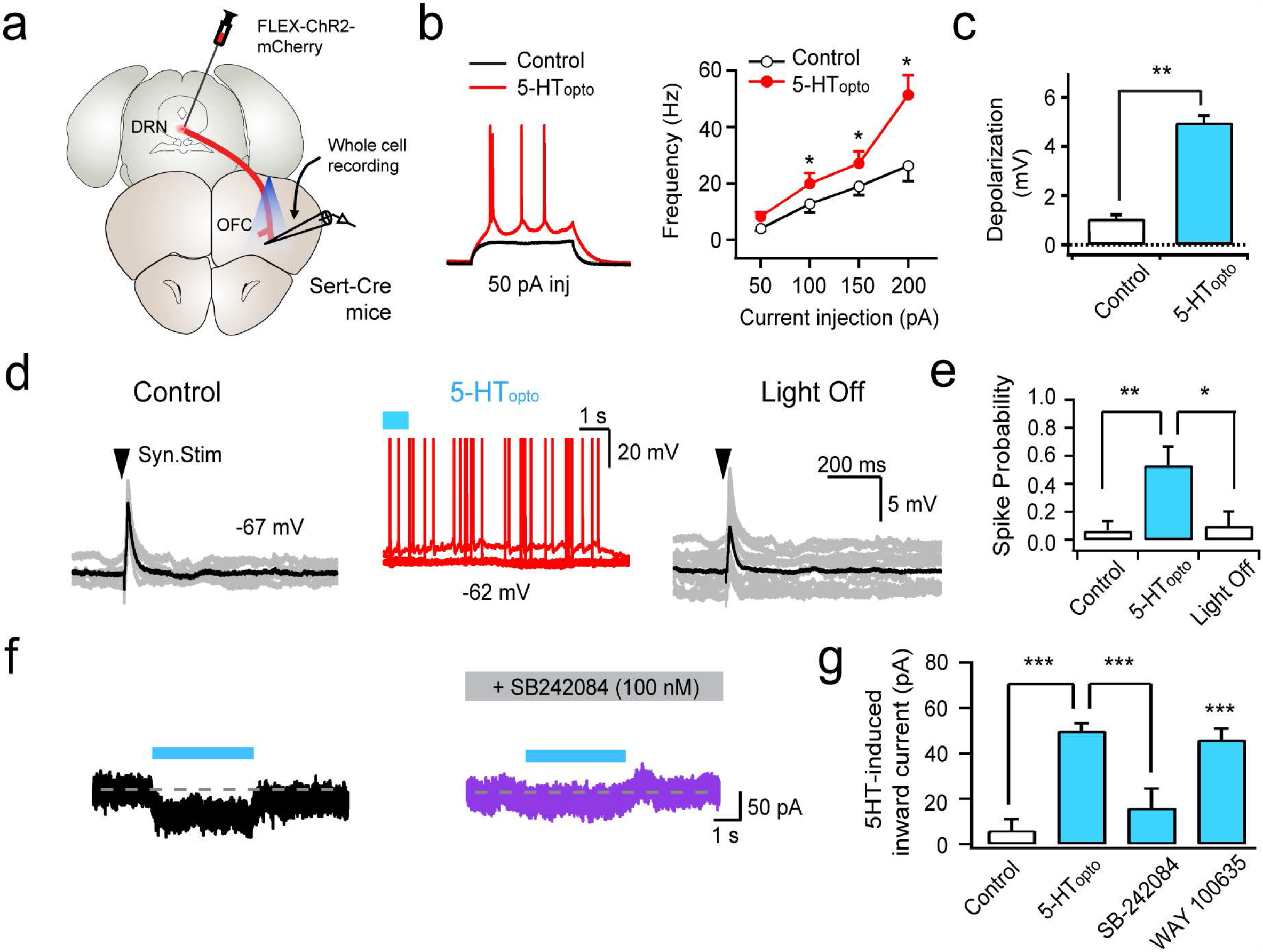
Serotonin increases the excitability of OFC neurons. **a**, Illustration of virus injection and whole-cell recording. **b**, Sample traces (left) and average frequency of action potentials (right) before and after 5-HT release. A brief current was injected to depolarize neurons. **c**, Average membrane potential changes by 5-HT (Control: 1.1 ± 0.5 mV; 5-HT_opto_: 4.2 ± 0.7 mV, n = 6, p < 0.01). **d**, Representative traces of facilitated E-S coupling by 5-HT. **e**, An average graph of spike probability changes (Control: 0.07 ± 0.04; 5-HT_opto_: 0.53 ± 0.15, n = 6, p < 0.01 compare to control; Light Off: 0.10 ± 0.07 mV, n = 6, p < 0.05 compare to 5-HT_opto_). **f**, Representative traces of depolarization triggered by blue light illumination. **g**, A summary graph showing serotonin-induced inward current and its blockade by SB-242084 (100 nM), but not WAY 100635 (1 mM) (Control: 6.1 ± 1.9 pA, n = 7; 5-HT_opto_: 50.2 ± 3.1 pA, n = 21, p < 0.005 compare to control; SB-242084: 16.0 ± 8.5 pA, n = 4, p < 0.005 compare to 5-HT_opto_; WAY 100635: 46.1 ± 4.6 pA, n = 8, p < 0.005 compare to control).

**Fig. 5.**
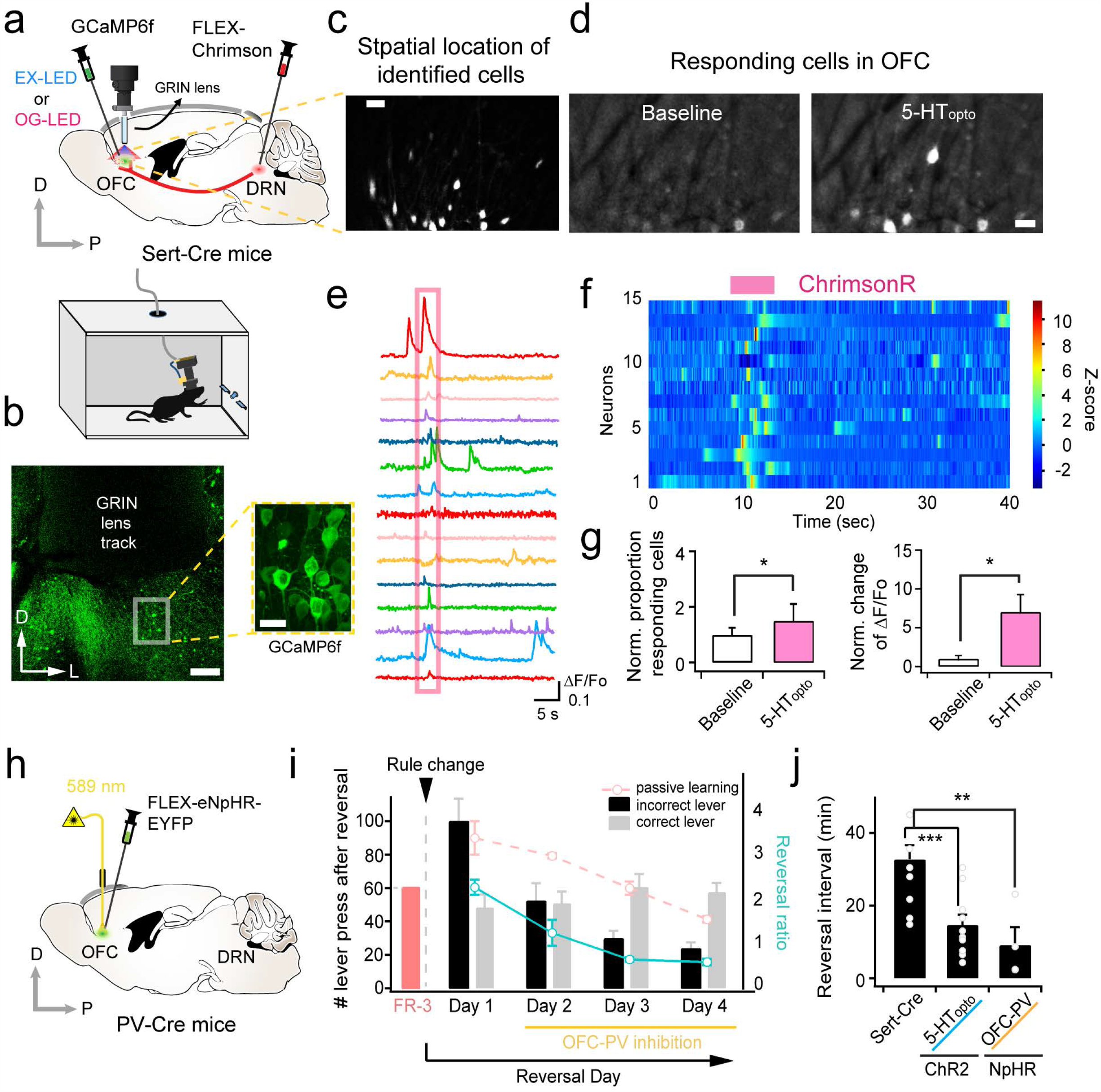
The role of serotonergic inputs to OFC *in vivo* and the impact of neuronal excitability on reversal learning. **a**, A schematic figure for Ca^2+^ imaging through GRIN lens affixed to a miniature microscope. **b**, Verification of correct targeting of virus injection and GRIN lens implantation. Confocal images were taken after fixation. Scale bars, 150 μm, 15 μm. **c**, A sample stack image of GCaMP6f signals in OFC neurons. Scale bar, 20 μm. **d**, Representative images showing Ca^2+^ fluorescence before and after 5-HT release obtained by miniscope imaging. Scale bar, 20 μm. **e**, Ca^2+^ transients from individual neurons in the OFC are aligned with different colors. A box indicates a period of optogenetic light illumination. **f**, A color map of Ca^2+^ signal changes. **g**, Summary bar graphs for the change in proportion of responding cells (left) or Ca^2+^ transient (right) induced by 5-HT_opto_ (Baseline: 1.0 ± 0.2; 5-HT_opto_: 1.5 ± 0.6, n = 13, p < 0.05); (Baseline: 1.0 ± 0.4; 5-HT_opto_: 7.0 ± 2.2, n = 13, p < 0.05). **h**, Illustration of virus injection scheme to selectively target PV-positive interneurons. **i**, A summary plot of reversal learning enhancement by PV-neuron inhibition (1-way RM-ANOVA, day: *F*_(1.8,10.7)_ = 22.6, p < 0.005; 5-HT_opto_ effect: *F*_(1,6)_ = 109.8, p < 0.005). **j**, Differences of reversal intervals at different conditions (Sert-Cre: 32.7 ± 3.9 min, n = 10; 5-HT_opto_: 14.7 ± 2.8 min, n = 10, p < 0.005 compare to Sert-Cre; OFC-PV^NpHR^: 9.2 ± 4.9 min, n = 4, p < 0.01 compare to Sert-Cre). ** and *** indicate p < 0.01 and p < 0.005, respectively. Error bars indicate s.e.m.

To examine whether 5-HT increases OFC neuronal excitability *in vivo*, we monitored Ca^2+^ signals from individual neurons through a miniaturized endoscope (miniscope). After injecting AAV-hSyn-GCaMP6s into the OFC, we implanted a gradient-index (GRIN) lens. Then, we attached the miniscope to monitor neuronal activity in behaving animals (Fig. 5a,c). Post-hoc analysis confirmed the precise location of virus injection and GRIN lens implantation in the OFC (Fig. 5b). To elicit firing from 5-HT neurons, we also injected AAV-Flex-ChrimsonR-mCherry^40^ into the DRN of Sert-Cre mice. Once a focal plane of GCaMP6f signals was identified, 620 nm light-emitting diode (LED) was delivered during behavior. We found that the magnitude of Ca^2+^ transients was increased upon illumination, implying more cell firing (Fig. 5d-g). Thus, these results indicate that the primary action of 5-HT in the OFC is to increase the excitability of neurons.

### Optogenetically Inhibiting PV-Interneurons in the OFC Promotes Reversal Learning

If the excitability change is a key cellular mechanism that accounts for RL, we may be able to control RL by manipulating the excitability of OFC neurons without activating serotonergic neurons. In the mammalian cortex, many investigators have manipulated interneuron activity as means of excitability changes^41-44^. Parvalbumin (PV)-positive interneuron has been shown to control the activity of pyramidal neurons in OFC^45,46^. Therefore, we inhibited the activity of PV interneurons to increase the excitability of pyramidal neurons. PV-Cre mice were trained, and AAV expressing Cre-dependent halorhodopsin (NpHR) was injected to the OFC. 589 nm light significantly improved RL, demonstrating enhancing excitability in OFC is sufficient to facilitate the process of disengaging with previous learning and ultimately improve RL (Fig. 5h-j).

A previous study reported that 5-HT neurons fire by both positive and negative prediction errors ^29^. This firing pattern and one of our main findings, excitability change, may explain how flexibility is encoded in individual neurons. For example, learning-related synaptic inputs transmit meaningful information by evoking action potentials, but when excitability increases, cells tend to fire indiscriminately. Thus, increased excitability eliminates the difference between learning-related inputs and non-relevant ones. This could be a cellular mechanism by which neuronal firing associated with previous learning gets weakened. Simultaneously, Ca^2+^ influx through N-methyl-D-aspartate (NMDA) receptors is enhanced due to depolarization^47,48^. These two mechanisms— increased excitability and increased Ca^2+^ entry—may be essential cellular features of reversal learning.

### Serotonin Facilitates Structural Plasticity at Single Dendritic Spine

To test this hypothesis, we examined local Ca^2+^ dynamics and synaptic plasticity mechanisms at single dendritic spine. First, we optimized the condition where we could obtain a well-isolated dendritic spine image (Fig. 6a-c). To monitor Ca^2+^ transients in dendritic spines, we injected a mixture of CaMKII-Cre and Flex-GCaMP6s viruses (Fig. 6d). After we made acute brain slices, 4-methoxy-7-nitroindolinyl-glutamate (MNI-Glu) was perfused in a slice chamber (2.5 mM) and illuminated ∼15 mW of 720 nm light 0.5 μm away from a single dendritic spine. Reliable Ca^2+^ transients were triggered by a two-photon light (Fig. 6e).

**Fig. 6.**
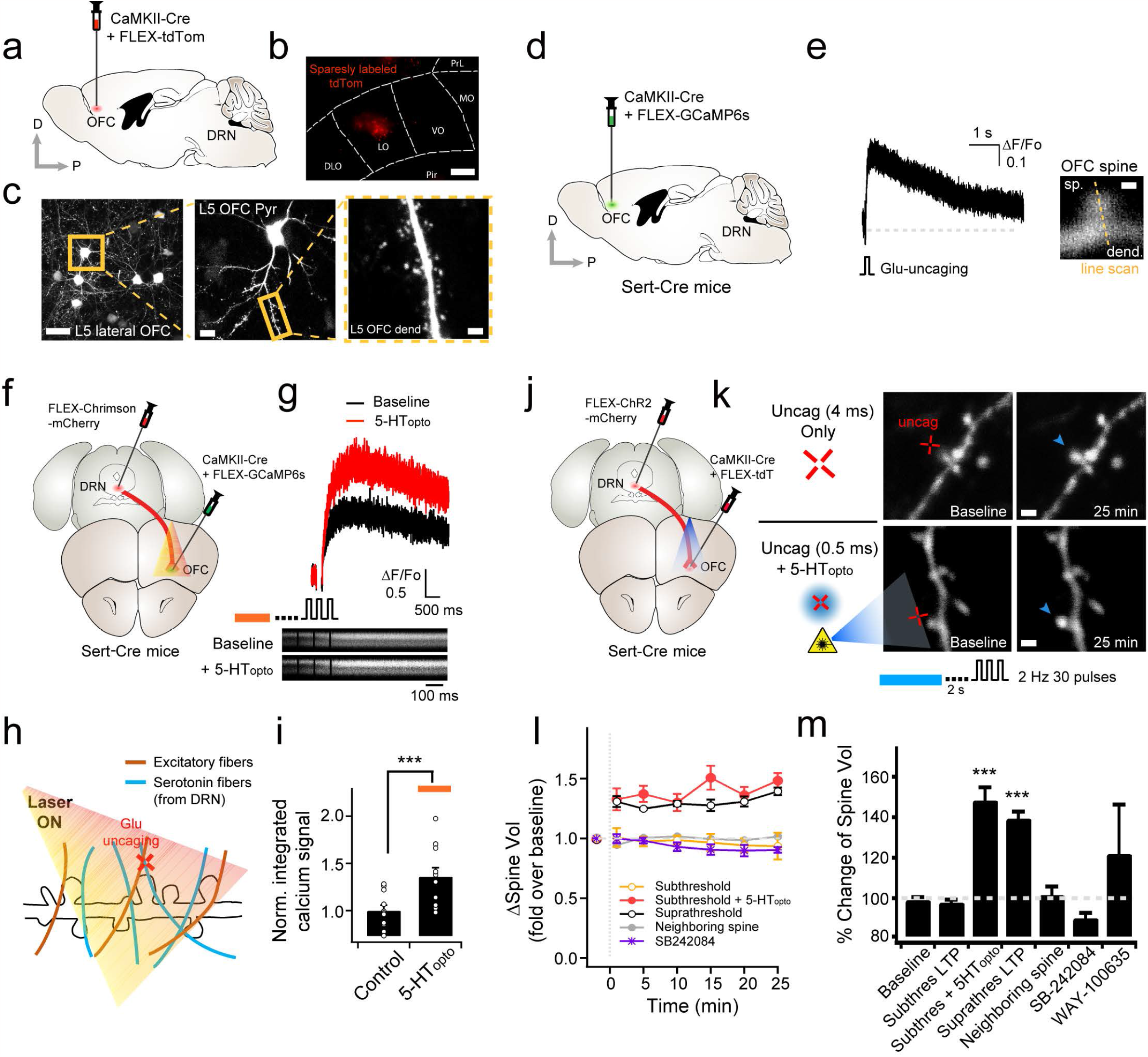
Two-photon Ca^2+^ imaging and structural plasticity from single dendritic spine. **a**, A virus injection scheme for sparse labeling. **b**, Representative image of tdTomato expression under low magnification. Scale bar, 200 μm. **c**, OFC pyramidal neurons were sparsely labeled to identify well-isolated dendrites and dendritic spines under 2P microscopy. Scale bars indicate 30 μm, 15 μm, and 5 μm, respectively. **d**, A virus injection scheme for Ca^2+^ imaging from the single dendritic spine. **e**, Excitatory postsynaptic current was evoked by two-photon glutamate uncaging (left). Ca^2+^ rise evoked by glutamate uncaging was visualized by line scanning across spine head and dendrite (right). **f**, A cartoon for virus injection and optogenetic stimulation. **g**, Sample traces of Ca^2+^ transients before and after 5-HT_opto_. **h**, Illustration of glutamate uncaging experiments at single spine resolution. **i**, Summary graph of integrated Ca^2+^ signals (Control: 1.0 ± 0.1; 5-HT_opto_: 1.4 ± 0.1, n = 11, p < 0.005). **j**, A virus injection scheme for testing structural plasticity. **k**, Representative images of structural plasticity triggered by 4 ms uncaging (top) and 0.5 ms uncaging together with 5-HT_opto_. Cross indicates the uncaging spot and arrowheads point enlarged spines. Scale bars indicate 2 μm. **l**, A time course graph of spine volume changes at different conditions. **m**, A summary plot of spine volume changes measured at 25 min after the LTP induction (Baseline: 98.8 ± 1.2%, n = 6; Subthres: 97.6 ± 5.2%, n = 4; Subthres + 5-HT_opto_: 148.6 ± 4.7%, n = 10, p < 0.005 compare to baseline; Suprathres LTP: 139.4 ± 3.2%, n = 4, p < 0.005 compare to baseline; Neighboring spine: 103.6 ± 6.7%, n = 5; SB-242084: 89.4 ± 4.1%,n = 8; WAY-100635: 123.0 ± 18.8%, n = 5). For all graphs, *** indicates p < 0.005, and Error bars indicate s.e.m.

To determine whether 5-HT alters the level of Ca^2+^ rise, we also injected AAV-Flex-ChrimsonR-mCherry into the DRN (Fig. 6f). When serotonin fibers were activated by 5-HT_opto_, the same glutamate uncaging power evoked a higher magnitude of Ca^2+^ rise, suggesting that 5-HT boosted Ca^2+^ influx (Fig. 6g-i). Elevated Ca^2+^ could lower the threshold of synaptic plasticity. We tested this possibility by testing activity-dependent spine growth (or enlargement), which have been known for a structural correlate of long-term potentiation (LTP)^49-53^. In particular, we used two different plasticity-induction protocols (suprathreshold vs subthreshold) by adjusting the duration of laser pulses of glutamate uncaging (4 ms vs 0.5 ms, respectively). 4 ms duration of laser pulses (30 pulses at 2 Hz) sufficiently triggered structural LTP (Fig. 6k-m). On the other hand, the subthreshold protocol (0.5 ms duration) was insufficient to triggering spine enlargement (Supplementary Fig. 6a,b), but when combined with blue light (to release 5-HT), the same uncaging protocol induced LTP (Fig. 6j-m). The LTP was blocked by a 5-HT_2C_ antagonist, SB242084 (Supplementary Fig. 6b), but not by a 5-HT_1A_ antagonist, WAY 100635 (Fig. 6l,m), indicating that the induction of LTP was facilitated by 5-HT. This metaplastic change by lowering the threshold of the LTP induction may account for capability of flexible updating of information.

## Discussion

In this study, we examined cellular- and circuit mechanisms by which 5-HT promotes RL. We targeted a genetically defined serotonergic neuronal group in the DRN and its projection to the OFC. Optogenetic manipulation of this circuit showed bidirectional up- and down-regulation of RL, indicating that the DRN-OFC circuits are necessary and sufficient of RL. 5-HT increased OFC neuronal excitability mediated through the activation of 5-HT_2C_ receptors. 5-HT also lowered the threshold of synaptic plasticity by allowing higher Ca^2+^ influx, rendering neuronal states more sensitive to the incoming inputs. Capability of switching choice behaviors was tightly linked to the state change of OFC neurons, implying that a general feature of cognitive flexibility may be represented as a form of internal state-dependent plasticity rather than a specific cell types or circuits. These serotonergic action-induced cellular and circuit changes in the OFC provide a new aspect of cellular features underlying cognitive flexibility.

The behavioral essence of cognitive flexibility is to maximize rewards in a changing environments. When the outcome of a decision constantly violate expectations, animals become alert and execute different behavioral strategies. To explain these behavioral characteristics, previous studies suggested that the primary function of OFC is a response inhibition of the previous learned behavior (*Reviewed in* ^8^). This hypothesis was supported by deficits of RL upon OFC area damage^11,17,54-56^. However, other studies also showed that animals with OFC lesions were normally able to hold decision to choose rewards that have been associated with devaluation^10,11,57,58^. Recent study also showed evidence of distinct segregated OFC pathways mediating the specific action-value updating and maintenance^59^. Thus, OFC neurons seem to have multifaceted functions and it is difficult to explain cognitive flexibility by one unifying mechanism. Further studies with cellular level analysis may be required for better understanding.

In this study, we dissected DRN-OFC circuits in an individual cell resolution. Looking at individual neuronal activity in the OFC uncovered that the major changes caused by 5-HT was the membrane potential depolarization. These findings were somewhat different from the previously suggested inhibitory actions of OFC. Instead, increased excitability indicates that a key mechanism of cognitive flexibility may be a transitory network state changes in which all incoming inputs become indistinguishable. This idea was confirmed by directly controlling network excitability without manipulating 5-HT levels in the OFC (Fig. 5). Another recent report also monitored RL after blocking PV neuron activities in the OFC^45^. Blockade of PV interneurons should increase the excitability of overall network activity, but they found that the opposite results, a deficit of RL, rather than an improved RL. However, it is interesting to note that they used prolonged light illumination protocol to inhibit PV interneurons during the entire sessions. In this case, increased global OFC network excitability would be sustained for long time, which may cause severe imbalance of excitatory and inhibitory network. This condition will cause the impairment of behavioral flexibility rather than the improvement.

Increase in excitability has also been proposed as a cellular mechanism of enhancing learning and allocating new memories^60,61^. Consistent with this idea, another key finding of our study is that excitability change facilitated the induction of synaptic plasticity. This metaplasticity was mediated by the activation of 5-HT_2C_ receptors. 5-HT_2_ receptors have been shown to produce neuronal depolarization and increased firing rates via multiple Gq protein-dependent manner, including opening of non-specific cation channels^62^. Our results are consistent with an effect of 5-HT action on the depolarization of the membrane potential and enhanced spike probability in OFC neurons (Fig. 4). Neuromodulatory 5-HT inputs acting on Gq-coupled receptors may show similar effects on LTP^63^. We found that subthreshold input-specific two-photon glutamate uncaging was able to make spine enlargement when combined with 5-HT release (Fig. 6). To our knowledge, the present study is among the first to describe the role of serotonergic inputs to the OFC on RL and identify potential underlying neural mechanism that regulates the modification of brain circuits at single cell and synapse resolution.

These cognitive and biological advances notwithstanding, our work is also implicated in clinical questions. Previous studies suggest that OFC is strongly associated with behavioral flexibility in the context of RL, whose abnormalities were consistently present in obsessive-compulsive disorder (OCD) patients^1,64,65^. Moreover, it has been hypothesized that serotonergic dysfunction is central to OCD pathophysiology^66^. Thus, impaired 5-HT signaling in OFC could impact the ability to rapidly adapt to new information, contributing to cognitive flexibility in OCD. We speculate that the understanding serotonergic signaling pathways may provide an effective means for modulating cognitive flexibility in patients affected by these disorders.

## Methods

### Animals

Experimental subjects were from 4 to 12 weeks old Sert-Cre (strain name B6.129(Cg)-Slc6a4tm1(cre)Xz/J) or C57BL6J mice (both from Jackson Laboratory, Bar Harbor, ME, USA). Similar number and ages of both male and female mice were randomly chosen for experiments. All mice were individually housed in a 12-h dark-light reverse cycle. All the behavior tests were performed during the dark cycle period. For the behavioral training, water was restricted to 1 ml per day for 3 days prior to training. Mice had free access to food ad libitum. All experimental procedures and protocols were conducted with the approval of the Max Planck Florida Institute for Neuroscience (MPFI) Institutional Animal Care and Use Committee (IACUC), Johns Hopkins University IACUC and National Institutes of Health (NIH) guidelines.

### Animal surgery and viral vectors

The general surgical procedure has been previously described^67^. Surgeries were performed on 4–6 week-old mice. A cocktail of ketamine (0.1 mg/g) and xylazine (0.01 mg/g) (Sigma-Aldrich) was used to anesthetize mice via intraperitoneal injection.

### Surgery for in vivo opto/chemo-genetics

#### In vivo optogenetic activation of cell body

In order to target the dorsal raphe nucleus (DRN) or orbitofrontal cortex (OFC), 300–400 nl of viral solution (pAAV1-EF1a-DIO-hChR2(H134R)-mCherry-WPRE-hGHn or AAV1-CAG-hChR2(H134R)-mCherry-WPRE-SV40) was injected into the DRN (Coordinates: AP, –4.7 mm; ML, 0 mm from bregma, and DV –2.85 mm from the brain surface; 32 degree tilted injection from rostral to caudal) or lateral part of OFC (Coordinates: AP, +2.65 mm; ML, ±1.65 mm from bregma, and DV –2 mm from the brain surface) through a glass micropipette (speed: 100∼200 nl/min). Following virus injection, the micropipette was held for 3 minutes to prevent backflow of viral solutions. Optical fiber (low OH, 200 μm core, 0.37 NA; Cat# BFL37-2000, Thorlabs) was fabricated as previously described^67^, and the optic fiber was placed in a ceramic ferrule and inserted into the brain region via craniotomy. The ceramic ferrule was supported with dental cement (C&B-Metabond, Parkell inc, Edgewood, NY). The tip of the fiber was positioned 150–200 μm above the target viral injection site in either DRN or OFC area. The following optic fiber implantation coordinates targeting the DRN or lateral OFC were used: AP, –4.7 mm; ML, 0 mm; and DV –2.65∼2.70 mm (32° angle using the same hole) or AP, +2.65 mm; ML, ±1.65 mm; and DV –1.8 mm, respectively. All values are given relative to the bregma.

#### In vivo manipulation of neuronal pathways from the DRN to the OFC

In order to manipulate the specific activity of OFC-projecting DRN neurons from Sert-Cre mice, 750 nl of viral solution (pAAVrg-EF1a-DIO-hChR2(H134R)-EYFP or pAAVrg-hSyn-DIO-hM4D (Gi)-mCherry) was injected into the OFC (see above virus injection coordination). After injection, an optic fiber was implanted in the DRN (see above fiber implant coordination). No optic fiber was implanted for chemogenetic inhibition experiments.

#### In vivo optogenetic activation of 5-HT axon terminals at OFC

In order to activate DRN 5-HT neuronal axon terminals in the OFC, 300–400 nl of viral solution (AAV1-EF1-DIO-hChR2(H134R)-mCherry-WPRE-hGHn) was injected to the DRN of the Sert-Cre mice (see above virus injection coordination). Following virus injection, an optical fiber was implanted in to OFC (see above fiber implant coordination).

### Surgery for retrograde viral tracings

To map out the circuit projection from the DRN to the OFC, AAVrg-pCAG-FLEX-tdTomato-WPRE (750 nl from each hemisphere) was delivered into the OFC and pENN-AAV1-CMV-PI-Cre-rBG (750 nl) was injected into the DRN of wild-type mice and allowed 2–3 weeks for retrograde transport. To target serotonergic neurons, 750 nl of AAVrg-pCAG-FLEX-tdTomato-WPRE was injected to the OFC of Sert-Cre mice, such that virus can retrogradely transport to the DRN and express Cre specifically in OFC-projecting DRN 5-HT neurons.

### Surgery for miniature microendoscope imaging

The surgeries consist of three steps; 1) Virus injection expressing opsins into the DRN (–5 weeks), 2) Cortical tissue aspiration followed by GRIN lens implantation after GCaMP injection in the OFC (–3 weeks), and 3) Baseplating procedure (0 week).

1) For the simultaneous imaging of OFC neurons and optogenetic stimulation of DRN 5-HT neurons, pAAV-Syn-FLEX-ChrimsonR-tdT (350 nl) was injected into the DRN of Sert-Cre mice (similar with aforementioned procedure) using the glass micropipettes (tip size 10–20 μm diameter, Braubrand) at the following coordinates from bregma: DRN (Coordinates: AP, –4.7 mm; ML, 0 mm from bregma, and DV –2.85 mm; 32 degree tilted injection from rostral to caudal).

2) Two weeks after opsin virus injection, the mice were anesthetized using isoflurane (4–5% for induction, 1.5–2.0% for maintenance; Patterson Vet Supply, Saint Paul, MN) and fixed them onto a stereotaxtic frame (Kopf instruments, Tujunga, CA, USA) for cortical aspiration, and GCaMP injection followed by GRIN lens implantation (GRIN lens, 1×4 mm, PN: 1050-002202, Inscopix, Palo Alto, CA) into the OFC. The head was shaved using hair remover lotion (Nair, Church & Dwight Co, Inc; Princeton, NJ) and petrolatum ointment (Puralube Vet Ophthalmic Ointment) was applied to both eyes to relieve dry eyes and surgical region was sterilized with alcohol and 10% betadine solution (Purdue product LP, Stamford, CT, USA). During surgery, core body temperature was maintained constantly at 37°C using a homeothermic blanket with flexible probe (Harvard Apparatus, Holliston, MA, USA). After checking the level of anesthesia, head skin and periosteum were carefully removed by a sharp surgical scissor and scalpel within aseptic surgical conditions. A craniotomy above the OFC (coordinates from bregma: AP, +2.65 mm; ML, +1.25 mm from bregma; this revised ML coordinate also centers over lateral part of OFC; reference of mouse brain atlas^68^ for GRIN lens implantation, was performed using a trephine (1.5 mm diameter, Fine Science Tools, Foster City, CA) with hand-held drill (Fordom Electric Co., Bethel, CT) and dura was carefully removed under the surgical microscope. 150 μm of tissue was aspirated using a 27-guage blunt needles attached to a vacuum pump to allow for an entry path for the GRIN lens. Saline was continuously applied during the aspiration to avoid drying of the tissue. Bleeding was controlled by sterile saline-soaked gelfoam (Pfizer, New York, NY) and cotton swab. Following aspiration, pAAV-Syn-GCaMP6f-WPRE-SV40 (350 nl) was unilaterally injected into the OFC (Coordinates: AP, +2.65 mm; ML, +1.25 mm from bregma, and DV –2 mm). Immediately after injection, mice were implanted with a cuffed 4 mm long GRIN lens (final target: AP, +2.65 mm; ML, +1.25 mm from bregma, and DV –1.8 mm) to enable optical access to the OFC and stimulation of axonal terminals of DRN 5-HT neurons that project to the OFC. During implantation, lenses were held in place and slowly lowered using a Pro View kit (1050-002334, Inscopix, Palo Alto, CA). The lens lowering speed was 300 μm/min for the first 1.5 mm (intended depth for aspiration), and then 100 μm/min for the remaining depth. The edges of the craniotomy were sealed with Kwik-Sil (World Precision Instrument, Sarasota, FL). Metabond (Parkwell, Edgewood, NY) was then applied around the lens to secure it in place. PCR tube sealed to head cap with Kwik-Sil was used to cover and protect the lens. The gap between the GRIN lens and the skull was covered with Metabond (Parkell, Edgewood, NY). A silicon elastomer (Kwik-Sil, World Precision Instruments) was applied to the top of the protective lens cuff to protect the top of the lens from debris or damage.

3) Three weeks following the lens implantation, the mouse was attached to a stereotaxic frame again to support the integrated microscope on top of the mouse’s head using a baseplate (1050-002192, Inscopix, Palo Alto, CA). The Kwik-Sil was detached from the lens cuff and the top of the implanted lens. The lens and cuff objective of the microscope were aligned parallel to each other. Using the stereotaxic arm, the microscope gripper was lowered until the field of view was positioned at the focal plane, as determined by landmarks, such as blood vessels. Once appropriate Ca^2+^ signal with nVoke2 (Inscopix, Palo Alto, CA) were detected, a baseplate was attached using Metabond to support a miniature microscope. A baseplate cover (BPC-2, Inscopix, Palo Alto, CA) was then secured on the baseplate to protect the lens prior to imaging sessions.

### Reversal learning task

Mice were placed in the reverse light cycle room and received a hand care for 5 min every day for 5-6 days before the lever-press training started. Mice were water-restricted for 3–4 days. Behavioral training procedure is comprised of two phases. All behavioral training and testing were performed in a standard mouse operant chamber (Med-Associates, St. Albans, VT) placed in a sound attenuating cubicle (ENV-022MD, 22 cm × 15 cm × 16 cm). The first phase of training (Day 1–7) is to make mice learn that lever pressing is associated with water reward. In the continuous reinforcement (CRF) session, a mouse received a water reward provided from a retractable sipper tube extended into the chamber after each left- or right-lever press. When mice finished getting five and fifteen rewards (CRF 5Rs and CRF 15Rs), they were moved to the fixed ratio (FR) schedule, where mice had to press a lever two or three times consecutively regardless of left or right in order to get a water reward. This FR-2 or normal FR-3 (without reversal) session lasted 45 min or until mice received twenty rewards. The mice performed for 2 consecutive days (on day 6–7). The second phase of training was the reversal learning task period (on day 8–12). The training started with FR-3 session, but only one side of lever was active. During this period, mice had to press an active lever (either the left or the right lever as chosen in the previous day) three consecutive times with a restriction of 20 sec time-out between presses. No cue or stimulus was associated with the active lever. After 9 rewards, the active lever became inactive and the inactive one became active. Reversal learning session was finished when an animal made 11 correct responses after the rule change or 45 minutes had elapsed. Reinforcers earned, along with active and inactive lever presses, were recorded during each session. The total amount of water was matched to 1 ml per day throughout the training periods by subtracting the amount of water given during the training sessions.

### Optogenetic activation of 5-HT neurons (5-HT_opto_) during the reversal learning training sessions

The AAV-infected mice were trained as described above. In order to activate cell bodies or axonal terminals of 5-HT neurons in the DRN, blue light (473 nm wavelength; 5–10 mW output power measured at the fiber tip with continuous light output) was illuminated 30 sec after mice successfully accomplished the first 9 rewards (the rule change time). 10 epochs of light were delivered at 0.1 Hz. Each epoch consisted of 5 trains of 10 pulses at 20 Hz with an interval of 1 sec. The laser (MBL-FN-473, Changchun New Industries Optoelectronics Technology, Jilin, China) was triggered by TTL output from MED-PC IV.

### Cannula implantation for local microinfusion

The naïve mice with AAV injection expressing retrograde inhibitory DREADD were trained with the lever-press operant task for 5 days. Two days before the normal FR-3 training session, mice went through the surgery for cannula implant. For intracranial drug infusion, a guide cannula (26 GA, Plastics One, Roanoke, VA) was implanted into the lateral part of the OFC bilaterally at the following coordinates: AP, +2.65 mm; ML, ±1.7 mm from bregma, and DV –1.5 mm from the brain surface. A 2 mm stainless steel dummy cannula (33GA, Plastics One) was also placed in the inside of each cannula. In order to make a precise drug infusion to the OFC area, the tip of cannula was positioned at 0.2 mm above the target site. After the surgery, the mice were allowed to recover for 5–7 days, then the mice were subjected to the behavioral chamber and allowed to test FR-2 or FR-3 schedule.

### Chemogenetic manipulation and intracranial microinfusion of Fluoxetine

During the 1^st^ day of reversal (on day 8), normal training procedure was executed. On the second day of reversal, CNO (1.875 mg/kg, i.p.) was injected 1 hour before the training started. On the following day, the day of fluoxetine injection (probe test), cannula (33GA, 1.8 mm projection for OFC, Plastics One) was inserted bilaterally, which were connected to a 5 ml Hamilton syringe via PE50 tubing back-filled with fluoxetine. Thirty minute before probe test (30 min after CNO treatment), fluoxetine was intracranially delivered with a volume of 58.7 nl/side over 2 min using a motorized single pressure syringe pump (World Precision Instruments, Sarasota, FL). After infusion, the cannula was left in place for an additional 5 min and then animals performed the reversal learning task.

### Measurement of 5-HT concentration in OFC area using *ex vivo* voltammetry

In order to measure 5-HT concentration in the OFC area, 500–600 nl of AAV1-EF1-DIO-hChR2-mCherry-WPRE-hGHn was injected into the DRN of Sert-Cre mice (see above virus injection coordination). 5–8 weeks following the virus injection, mice were anesthetized with isoflurane and decapitated to obtain OFC-containing slices (See electrophysiological recordings section). Fast-scan voltammetry recording was performed using carbon fiber electrodes (7 μm diameter, Tex # 795, Goodfellow, Coraopolis, PA). Carbon fibers were enclosed in pulled-glass capillaries (8250, A-M Systems, Sequim, WA) with back-filled epoxy (Epo-Tek 301, Epoxy Technology, Billerica, MA) and were cut to the size of 50-100 μm exposed length. Prior to use, electrodes were cleaned with bleach (4-6% Sodium Hypochlorite, Hawkins, Roseville, MN) for 5 min to promote greater sensitivity. Electrodes were filled with 3M KCl and connected to a headstage (CV-7B with 50 MΩ feedback resistor, Molecular Devices, San Jose, CA) via Ag/AgCl wire. Each electrode was pre-calibrated with fresh 50-100 nM 5-HT, which exhibited +680 mV oxidation peak. The potential of the carbon electrode was scanned at a rate of 1000 V/s^69^ according to a triangular voltage waveform (+0.2 to +1V vs. Ag/AgCl reference) from +0.2V holding potential with 100 ms sampling intervals using Multiclamp 700B amplifier (Molecular Devices). Data were acquired using Axon Digidata 1550B (Molecular Devices) at 10 kHz sampling rate with 4 kHz low-pass filter. Background current was recorded for 30 sec and averaged, subtracted digitally using Clampfit 10.6 software (Molecular Devices). 473 nm wavelength of blue light stimulation was applied to slices through water-immersion 40X objective coupled to a LED (X-Cite 120LEDBoost, Excelitas Technologies, Mississauga, ON, Canada) and TTL-controlled by a pulse stimulator (2100 isolated pulse stimulator, A-M Systems). Photoelectric currents on carbon fibers have been reported when optogenetically stimulated^70-72^. To minimize photoelectric current, shorter pulse duration (4 ms) was used ^72^ and electrode was inserted deeper^70^ into the slice, approximately 150 μm in depth. Light output power was estimated using an optical power meter (PM100D with S130C photodiode sensor, Thorlabs, Dachau, Germany). 100 Hz (500 pulses with 4 ms, 28 μW), 20 Hz (100 pulses with 4 ms, 140 nW), and 5 sec for a prolonged stimulation (1.8 mW) were measured and the ability of terminal stimulation to release 5-HT using ex vivo voltammetry was verified. Data were analyzed with OriginPro 2017 (OriginLab, Northampton, MA), and Prism 6 (GraphPad software, La Jolla, CA).

### Electrophysiological recordings

In order to test the direct effect of 5-HT on OFC neurons, 300– 400 nl of viral solution (AAV1-EF1-dFlox-hChR2-mCherry-WPRE-hGHn) was injected into the DRN of Sert-Cre mice. Details of the brain slice preparation is previously described^67^. Briefly, 5– 9 weeks after virus injection, mice were deeply anesthetized using isoflurane and the brain was quickly removed and chilled in an ice-cold high-magnesium cutting solution containing the following (in mM): 116 NaCl, 26 NaHCO_3_, 3.2 KCl, 0.5 CaCl_2_, 7 MgCl_2_, 1.25 NaH_2_PO_4_, 10 glucose, 2 Na-pyruvate, 3 ascorbate, with pH 7.4 adjusted by carbogen (95% O_2_, 5% CO_2_). Osmolarity was set to ∼300 mOsm. Coronal brain slices of 300 μm thickness were prepared by using a vibratome (Leica VT1000, Leica Biosystems, Buffalo Grove, IL) and incubated at 34°C for 30 min in the same solution, and thereafter maintained at room temperature for at least an hour. For experiments, we transferred the slice to a recording chamber superfused with artificial cerebrospinal fluid (ACSF) containing the followings (in mM): 124 NaCl, 26 NaHCO_3_, 3.2 KCl, 2.5 CaCl_2_, 1.3 MgCl_2_, 1.25 NaH_2_PO_4_, 10 glucose, bubbled with 95% O_2_ and 5% CO_2_.

Whole-cell current- or voltage-clamp recordings were carried out from layer V pyramidal neurons in the OFC while the recording chamber was continuously perfused with ACSF at 1–1.5 ml/min. The recordings were made using a MultiClamp 700B amplifier controlled by Clampex 10.2 and data were obtained by Digidata 1440A data acquisition system (Molecular Devices). The pipette solution contained (in mM): 125 K-gluconate, 5 KCl, 10 Na_2_-phosphocreatine, 4 Mg-ATP, 0.4 Na-GTP, 10 HEPES, 1 EGTA, 3 Na-ascorbate (pH = 7.25 with KOH, 295 mOsm). After forming whole-cell patch on the soma of a cortical neuron, we monitored E-S coupling, membrane potential changes and the number of action potentials before and after the optogenetic activation of 5-HT neurons (5-HT_opto_) under the current-clamp mode. ChR2 was activated by 470 nm wavelength LED (pE-100, CoolLED). Excitatory postsynaptic potentials (EPSPs) were evoked by stimulating superficial layers of cortex with 0.2-ms pulses (concentric bipolar electrodes, FHC). Under voltage-clamp modes, the magnitude of inward currents were measured upon 5 sec blue light.

### Optogenetic silencing of PV interneurons in the OFC

PV-Cre mice (Jackson Laboratory, Cat# 8069) were used for the experiments and treated for viral injection (pAAV-Ef1a-DIO-eNpHR-EYFP-WPRE-pA) and optic fiber implantation into the OFC using the procedures as described above. Animals were normally trained up to the first day of reversal learning. On the second day of reversal learning, yellow light (589 nm; 5–10 mW) was delivered 30 sec after the rule change through the bilateral optic fibers. 20 Hz, 10 pulses of light were shined 5 times at with an interval of 1 sec, and this train was repeated 10 times at 0.1 Hz. Over several days, the mice received the same light procedure during the reversal learning phase and of the performance of the animal during reversal learning was assessed. The number of lever press and the reversal ratio were recorded and analyzed.

### Integrated miniature microendoscope imaging

For simultaneous Ca^2+^ imaging and optogenetic stimulation, an integrated miniscope (nVoke2.0, Inscopix, Palo Alto, CA) was affixed to the baseplate described above (see the ‘surgery for miniature microendoscope imaging’ section). To monitor Ca^2+^ transients in OFC neurons, EX-LED (435–460 nm) was shined through the implanted GRIN lens, and to trigger 5-HT release from DRN 5-HT neurons, OG-LED (590–650 nm) was used. Prior to in vivo imaging experiments, Sert-Cre mice were injected with 350 nl of AAV1-Syn-GCaMP6f in the OFC and pAAV-Syn-FLEX-ChrimsonR-tdT in the DRN. Imaging data were acquired using nVoke Acquisition Software (version 2.0, Inscopix, Palo Alto, CA) at 20 frames/sec with a LED power of 40–60% (0.9–1.2 mW at the objective, 475 nm), analog gain of 2–4 and a field of view of 650×650 μm. For individual mice, the same imaging parameters were kept across repeated behavioral sessions.

In order to examine the physiological effect of 5-HT on the OFC neuronal excitability in fully awake mice, OG-LED (620 nm) was illuminated for 5 sec to activate ChrimsonR-expressing axon terminals of DRN 5-HT neurons while Ca^2+^ signals from the OFC neurons were recorded. Ca^2+^ imaging, optogenetic activation, and behavior were all timely-matched. All experiments were performed in an operant behavioral chamber (Med-Associates, St. Albans, VT). All image processing was performed using Mosaic Software (version 1.2, Inscopix, Palo Alto, CA). First, pre-processing was performed to remove irrelevant regions and lens borders. By using a spatial bandpassing gaussian filter, background was subtracted on each segment which was motion corrected by using the mean image (as the reference image) over time. Then, motion corrected segments were re-combined into a single movie and cropped to remove the edge effects. The reference image was computed to produce an F0 image, and the ΔF/F was calculated using baseline (with F0 = mean fluorescence of entire trace). After image processing, Ca^2+^ responses from individual cells were identified by Principal Component Analysis/Independent Component Analysis (PCA/ICA). Ca^2+^ transients were picked by evaluating the shape of the responses and their time scale. To create the heatmap of responses, the mean value (mean) and standard deviation (SD) of the GCaMP6f signals from each cell were used to acquire Z-score as follows: Z-score = {F(t)–mean}/SD, where F(t) is the GCaMP6f signal. The criteria for counting active cells was set by the fluorescence level ΔF/F 0.004%, and counted all cells exhibiting above that level. To measure the change of ΔF/F value in the presence of 5-HT_opto_, ΔF/F from GCaMP6-exrpessing OFC cellular region of interests (ROI) were averaged from the 2 OG-LED trials on time periods, and were normalized to a 30 s period of the baseline.

### Two-photon spine imaging and glutamate uncaging

A combined two-photon laser-scanning microscope imaging and two-photon photolysis of glutamate was performed with an upright microscope (Bruker, Inc) and a water-immersion objective lens (LUMPlanFL N, 60×, NA 1.0). Two mode-locked, femtosecond-pulse Ti:sapphire lasers (Mai Tai HP and Mai Tai DeepSee; Spectra-Physics) were used for uncaging and imaging at wavelengths of 720 and 920 nm, respectively. For optogenetic stimulation, light was delivered via FC/APC fiber (400 μm, 1m long, CNI, Jilin, China), coupled to either a blue or yellow laser (Changchun New Industries Optoelectronics Technology, Jilin, China) and was placed two centimeters above the surface of the cortical slice. The light intensity was adjusted to 5-10 mW measured at the tip of the fiber.

For spine imaging, sparse labeling was achieved by injecting a mixture (1:5) of AAV-CaMKII-Cre particle (60 nl) and AAV1-Flex-tdT (300 nl) to the OFC of Sert-Cre mice. This allowed us to detect well-isolated dendritic branches and dendritic spines labelled with tdTomato expression. 4– 7 weeks after virus injection, mice were deeply anesthetized using isoflurane and the brain was quickly removed to obtain acute slices (see method section for electrophysiological recording). The OFC neurons from the acute slice were imaged using two-photon microscopy with a pulsed Ti::sapphire laser. For each neuron, image stacks (512 × 512 pixels; 0.048 μm/pixel) with 1-μm z-steps were collected to cover the entire segment of apical dendrites. All images shown are maximum projections of 3D image stacks after applying a median filter (2 × 2) to the raw image data.

For glutamate uncaging, 4-methoxy-7-nitroindolinyl-glutamate (MNI-caged glutamate, Tocris) was dissolved in Mg2+-free ACSF (4 mM Ca^2+^, 0.5 μM TTX) to get the final concentration of 2.5 mM. Laser power was adjusted to 15 mW, which reliably triggered excitatory synaptic currents without causing phototoxicity^73^. Experiments were performed at room temperature (24–26 °C). To visualize Ca^2+^ signals, AAV-CaMKII-Cre and Cre-dependent GCaMP6s viruses were injected to the OFC. Dendritic spines in the oblique dendrites of layer 5 OFC pyramidal neurons were selected for Ca^2+^ imaging. 3 pulses of lasers (10 Hz) with 10 ms duration were delivered 0.5 μm away from the single dendritic spine, which triggered reliable Ca^2+^ rise from the spine head. To detect Ca2+ transients, we did line scan crossing the target dendritic spine head. The fluorescence changes were represented as ΔG/G_0_ = (G(t_peak_)−G_0_)/G_0_, where G(t_peak_) is the peak intensity of GCaMP6s fluorescence, and G0 is the average intensity before stimulation.

To compare Ca^2+^ transients in the 5-HTopto condition, we additionally injected pAAV-Syn-FLEX-ChrimsonR-tdT virus into the DRN. Ca^2+^ transients were compared before and after 5 sec of yellow light (MBL-F-589 nm, CNI, China) upon the same uncaging laser pulses.

### Structural plasticity of dendritic spines

To test the effect of 5-HT on spine structural plasticity, Sert-Cre mice were injected AAV-CaMKII-Cre (60 nl) + AAV1-Flex-tdT (300 nl) to the OFC and AAV1-EF1-DIO-hChR2-mCherry-WPRE-hGHn into the DRN (see above virus injection coordination).

Spine enlargement (also called structural LTP) was induced by repetitive glutamate uncaging (30 pulses, 2 Hz). We used two different plasticity-induction protocols (Suprathreshold vs subthreshold) by adjusting the duration of laser pulses of glutamate uncaging (4 ms vs 0.5 ms). Estimated spine volume was measured from bleed-through-corrected and background-subtracted red fluorescence intensities using the integrated pixel intensity value of a circular ROI which was placed over individual dendritic spine, as previously described^74^.

### Tissue fixation and acquisition of confocal imaging

General procedure for tissue fixation and confocal imaging was described previously ^67^. In brief, mice were sacrificed via deep anaesthetization with a mixture of ketamine and xylazine and then transcardially perfused with PBS (pH 7.4) followed by 4 % paraformaldehyde (PFA). The fixation procedure for GRIN lens implanted animals was similar, but the decapitated head was kept overnight in 4 % PFA at 4 °C with intact optical equipment and then brain was removed next day. Following post-fixation in 4 % PFA overnight at 4 °C, the mouse brain was embedded into 10 % gelatin solution at 50°C briefly and then embedded brains were kept in 4 °C for additional 30 min for solidification of gel. The gel was trimmed to a small cube around the brain and the cube was stored in 4 % PFA overnight. The gelatin cube was sectioned coronally with 100 μm thickness (except for sagittal sections tilted 12 degrees away from the midline in Fig. 1) using a vibratome (Leica VT1200). Imaging was performed using upright confocal laser-scanning microscope (LSM880, Zeiss, Germany) with 20×/0.8 M27 objective lens. Sagittal images from tilted sections were analyzed using a 10× objective lens. All the confocal images were analyzed by ImageJ (NIH).

### Viruses and pharmacology

All viruses were from Addgene unless noted otherwise. SB-242084, WAY-100635, Fluoxetine and Tetrodotoxin were purchased from Tocris.

### Statistical analysis

Data were analyzed using IgorPro (version 6.10A; WaveMetrics, Lake Oswego, OR). Statistical data were presented as means ± standard error of mean (SEM, denoted as error bars), and n indicates the number of cells or animals studied. The significance of differences between two experimental conditions was evaluated using Student’s t test, or Wilcoxon’s signed rank test for non-paired and paired data after testing normality using the Kolmogorov-Smirnov test. Behavioral parameters from reversal learning task were evaluated using one-way repeated-measures ANOVA or nonparametric Mann-Whitney U test. Difference were considered as significant when *P* < 0.05. n.s., no statistical significance; *, *p* < 0.05, **, *p* < 0.01, ***, *p* < 0.005.

## Acknowledgements

We thank members of the Kwon laboratory for helpful discussions; We thank W.C.Oh for helping two-photon uncaging. GCaMP6s virus was available from the GENIE Project and the Janelia Research Campus, especially V.Jayaraman, R.A.Kerr, D.S.Kim, L.L.Looger, and K.Svoboda. This work was supported by Max Planck Florida Institute (to H-B.K.), the National Institute of Health Grants DP1MH119428 (to H-B.K) and the National Institute of Health Award MH096972 (to RD.B.).

## Author contributions

J.H.H. and H-B.K. conceived and designed the study. J.H.H. performed all the experiments and analyzed data with the help of P.H. for animal surgery and behavioral training. H.I. and RD.B. performed a fast-scan cyclic voltammetry recording. J.H.H. and H-B.K. wrote the manuscript.

## Competing interests

The authors declare no competing interests.

## Notes

### Competing Interest Statement

The authors have declared no competing interest.

