## Supplementary Figures for "Serotonin in the orbitofrontal cortex enhances cognitive flexibility"

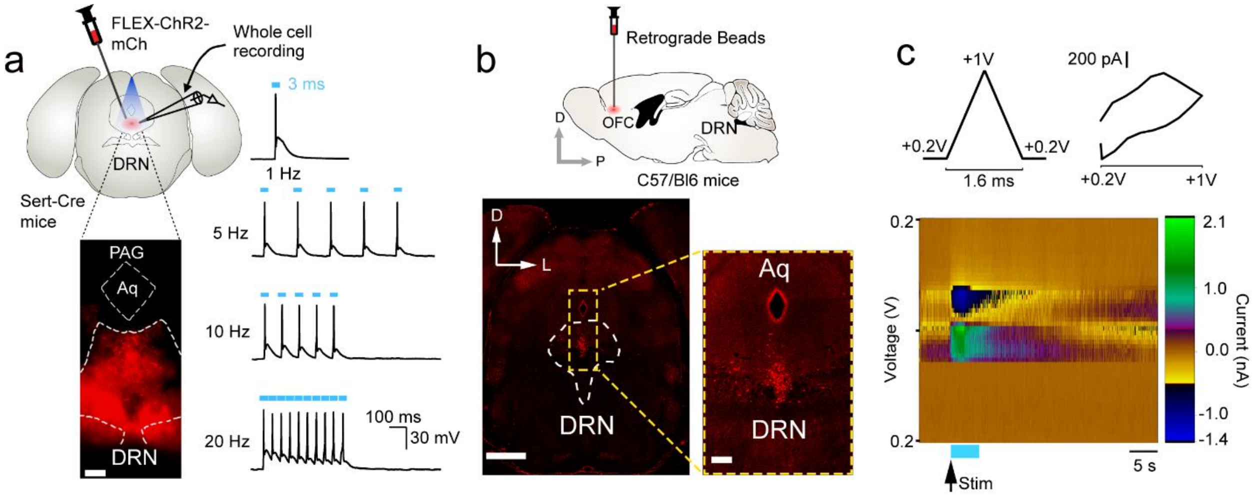


**Supplementary Fig. 1 Functional and anatomical verification of projections from the DRN to OFC. a**, Cre-dependent AAV expressing ChR2-mCherry was injected to Sert-Cre mice in order to selectively label serotonergic neurons. Confocal image showed precise targeting of virus injection to the DRN area (left). A whole-cell recording was made from cells expressing mCherry signals. Current-clamp recording was made to detect action potentials triggered by blue light. High frequency (20 Hz) of blue light reliably triggered action potentials (right). Scale bar, 100 µm. **b**, Retrograde beads were injected into the OFC in order to detect signals in the DRN (top). Signals from beads were visible in the DRN (bottom). Scale bars indicate 500 µm and 100 µm, respectively. **c**, Fast-scan cyclic voltammetry recording detected 5-HT release in the OFC by activating the axon terminals of serotonergic neurons. Triangular voltage wave (left) and voltage-current plot (right) were shown. Color representation of voltage and current change (bottom). Blue rectangle indicates blue light illumination.

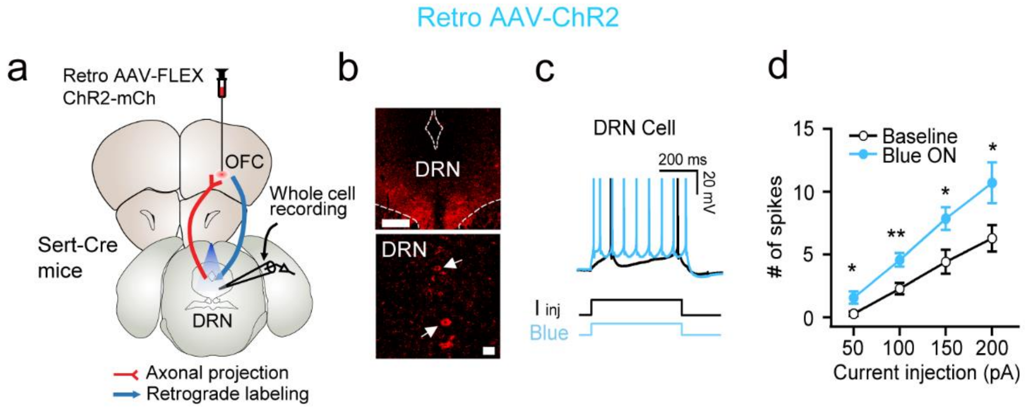


**Supplementary Fig. 2 Reliable action potential triggers by optogenetic stimulation of 5-HT neurons in the DRN. a**, A scheme for virus injection and whole-cell recording. Retrograde AAV expressing ChR2 was injected into the OFC and blue light was illuminated in the DRN. **b**, ChR2-mcherry signals were found in the DRN area indicating the existence of 5-HT neurons projecting to the OFC. Scale bars, 100 µm and 20 µm. **c**, ChR2 activation by blue light caused increased firing in 5-HT neurons. Sample traces and the current injection schematic were shown. **d**, Summary plot of action potential number (50 pA: 0.3 ± 0.2 for Baseline; 1.6 ± 0.5 for Blue ON, n = 7, $p < 0.05$ compare to baseline; 100 pA: 2.3 ± 0.5 for Baseline; 4.6 ± 0.5 for Blue ON, n = 7, $p < 0.01$ compare to baseline; 150 pA: 4.4 ± 0.9 for Baseline; 7.9 ± 0.9 for Blue ON, n = 7, $p < 0.05$ compare to baseline; 200 pA: 6.3 ± 1.1 for Baseline; 10.7 ± 1.6 for Blue ON, n = 7, $p < 0.05$ compare to baseline). * and ** indicate $p < 0.05$ and $p < 0.01$, respectively. Error bars indicate s.e.m.

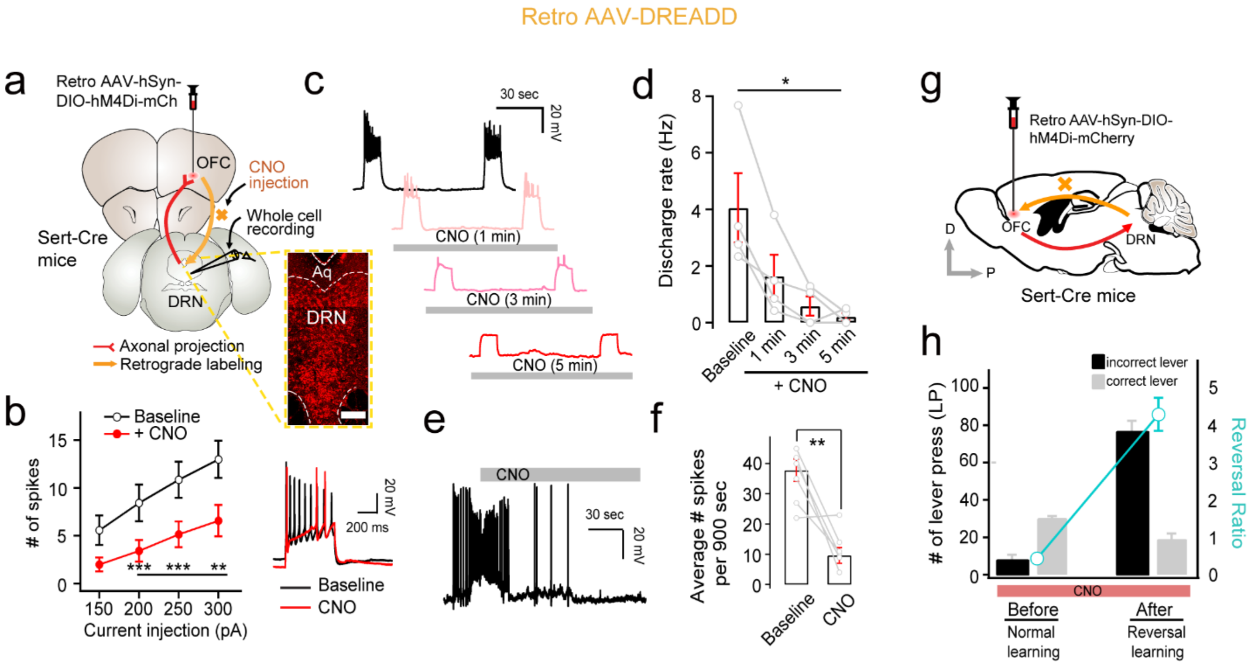


**Supplementary Fig. 3 Reduced cell firing and impaired reversal learning by CNO application.** **a**, A schematic figure illustrating virus injection and whole-cell recording sites. Whole-cell recording was made in neurons expressing hM4Di-mCherry. Scale bar, 100 µm. **b**, A summary graph of spike number with and without CNO (150 pA: 5.6 ± 1.5 for Baseline; 2.0 ± 0.7 for CNO, n = 7, n.s. compare to baseline; 200 pA: 8.4 ± 1.9 for Baseline; 3.4 ± 1.1 for CNO, n = 7, $p < 0.005$ compare to baseline; 250 pA: 10.9 ± 1.9 for Baseline; 5.1 ± 1.4 for CNO, n = 7, $p < 0.005$ compare to baseline; 300 pA: 13.0 ± 2.0 for Baseline; 6.6 ± 1.6 for CNO, n = 7, $p < 0.01$ compare to baseline). **c**, CNO efficiently reduced cell firing as time passed. **d**, The number of action potentials induced by current injection was reduced in the presence of CNO (Baseline: 4.0 ± 1.2; CNO: 1.6 ± 0.8 for 1 min, n = 4, $p < 0.05$ compare to baseline; 0.6 ± 0.3 for 3 min, n = 4, $p < 0.05$ compare to 1 min; 0.2 ± 0.1 for 5 min, n = 4, $p < 0.05$ compare to 3 min). **e**-**f**, Total number of action potentials during longer time scale (15 min) was compared before and after CNO application (Baseline: 35.2 ± 3.7; CNO: 11.8 ± 2.6, n = 6, $p < 0.01$ compare to baseline). **g**, Sagittal section view of virus injection and CNO effect. **h**, A summary plot showing the number of lever presses and reversal ratio. In the presence of CNO, normal lever pressing associative learning before reversal learning phase was not affected, but ability to switch the side of lever (reversal learning) was impaired. (Reversal ratio: 0.2 ± 0.1 for normal learning; 4.4 ± 0.5 for reversal learning, n = 8, $p < 0.005$). *, ** and *** indicate $p < 0.05$, $p < 0.01$ and $p < 0.005$, respectively. Error bars indicate s.e.m.

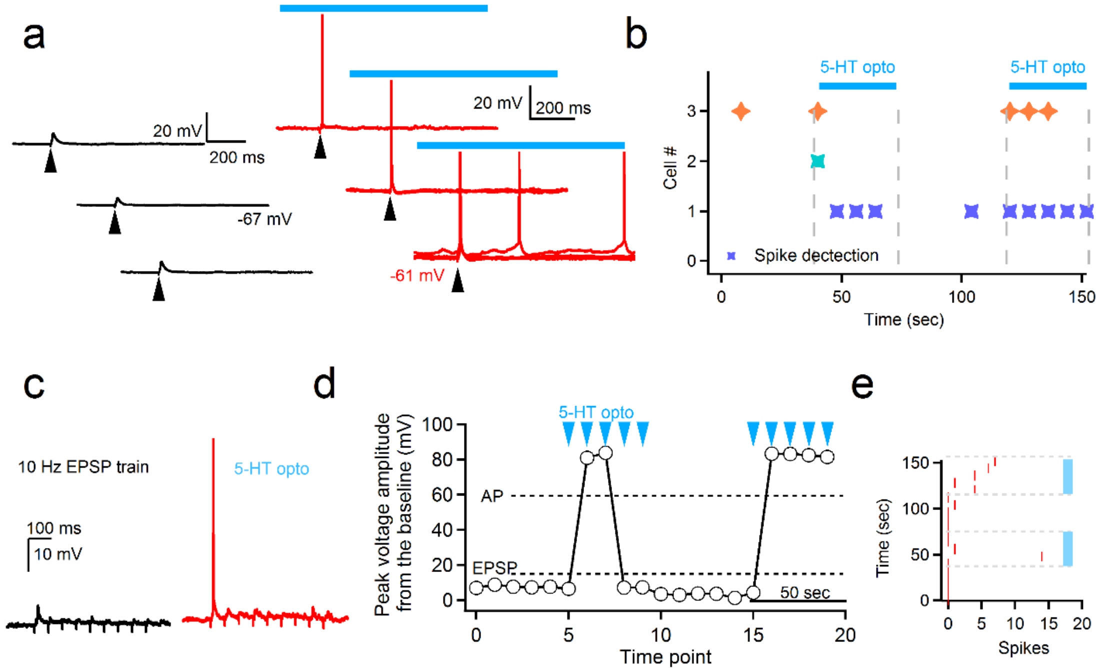


**Supplementary Fig. 4 Enhanced EPSP-to-Spike (E-S) coupling by 5-HT. a**, Example traces of excitatory postsynaptic potentials (left) in the OFC neuron and their conversion to action potentials (right) in the presence of 5-$HT_{opto}$. **b**, Time course plot demonstrating more spikes upon 5-HT release. **c**, Firing pattern change during a short train of stimulation. **d**-**e**, Repeatable and reversible switch from EPSPs to spikes depending on 5-HT release.

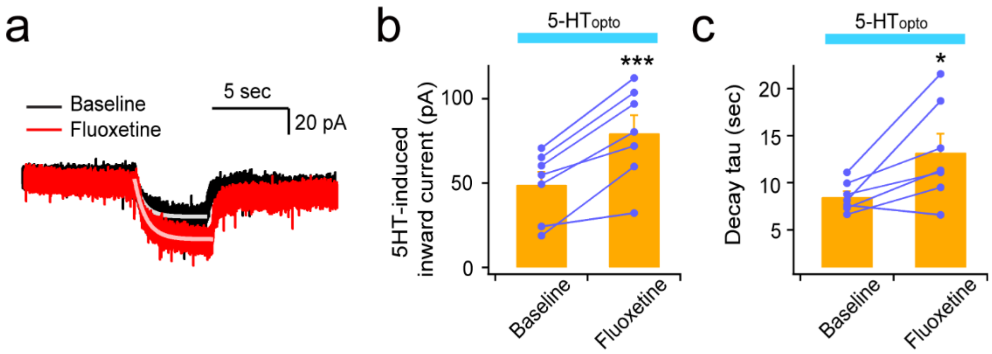


**Supplementary Fig. 5 5-HT-induced inward current is prolonged by fluoxetine. a**, Example trace of inward current induced by 5-$HT_{opto}$ in the absence (black) or presence (red) of 10 μM fluoxetine. **b**, Summary bar graph for the effect of fluoxetine on the peak amplitude of current induced by 5-$HT_{opto}$ (inward current: 49.2 ± 7.6 pA for Baseline; 79.7 ± 10.5 pA for Fluoxetine, $n = 7$, $p < 0.005$ compare to baseline). **c**, Summary bar graph for the effect of fluoxetine on the decay kinetics of currents induced by 5-$HT_{opto}$ (decay tau: 8.5 ± 0.6 sec for Baseline; 13.2 ± 2.0 sec for Fluoxetine, $n = 7$, $p < 0.05$ compare to baseline). * and *** indicates $p < 0.05$ and $p < 0.005$, respectively. Error bars indicate s.e.m.

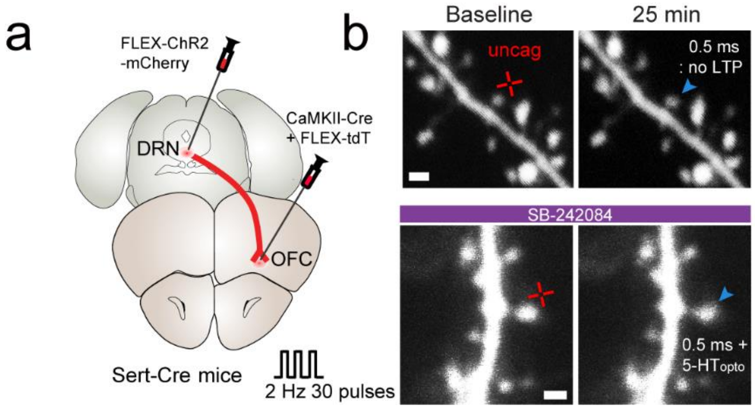


**Supplementary Fig. 6 Glutamate uncaging-induced spine enlargement test. a**, AAV-FLEX-ChR2-mCherry was injected into the DRN of Sert-Cre mice in order to control serotonin release. **b**, Representative images of no spine enlargement by 30 pulses of glutamate uncaging at 2 Hz with 0.5 ms duration (top). The same protocol combined with 5-HT release normally triggered structural plasticity, but was blocked in the presence of 5-$HT_{2C}$ antagonist, SB-242084 (bottom). Cross marks indicate uncaging spots. Scale bars indicate 2 µm.
